# Structural and functional evaluation of *de novo-*designed, two-component nanoparticle carriers for HIV Env trimer immunogens

**DOI:** 10.1101/2020.01.31.929273

**Authors:** Aleksandar Antanasijevic, George Ueda, Philip JM Brouwer, Jeffrey Copps, Deli Huang, Joel D Allen, Christopher A Cottrell, Anila Yasmeen, Leigh M Sewall, Ilja Bontjer, Thomas J Ketas, Hannah L Turner, Zachary T Berndsen, Per Johan Klasse, Max Crispin, David Nemazee, John P Moore, Rogier W Sanders, Neil P King, David Baker, Andrew B Ward

## Abstract

Two-component, self-assembling nanoparticles represent a versatile platform for multivalent presentation of viral antigens. Nanoparticles of different sizes and geometries can be designed and combined with appropriate antigens to fit the requirements of different immunization strategies. Here, we describe detailed antigenic, structural, and functional characterization of computationally designed tetrahedral, octahedral, and icosahedral nanoparticle immunogens displaying trimeric HIV envelope glycoprotein (Env) ectodomains. Env trimers, based on subtype A (BG505) or consensus group M (ConM) sequences and engineered with SOSIP stabilizing mutations, were fused to the underlying trimeric building block of each nanoparticle. Initial screening yielded one icosahedral and two tetrahedral nanoparticle candidates, capable of presenting twenty or four copies of the Env trimer. A number of analyses, including detailed structural characterization by cryo-EM, demonstrated that the nanoparticle immunogens possessed the intended structural and antigenic properties. Comparing the humoral responses elicited by ConM-SOSIP trimers presented on a two-component tetrahedral nanoparticle to the corresponding soluble protein revealed that multivalent presentation increased the proportion of the overall antibody response directed against autologous neutralizing Ab epitopes present on the ConM-SOSIP trimers.

**Author Summary:** Protein constructs based on soluble ectodomains of HIV glycoprotein (Env) trimers are the basis of many current HIV vaccine platforms. Multivalent antigen display is one strategy applied to improve the immunogenicity of different subunit vaccine candidates. Here, we describe and comprehensively evaluate a library of *de novo* designed, protein nanoparticles of different geometries for their ability to present trimeric Env antigens. We found three nanoparticle candidates that can stably incorporate model Env trimer on their surface while maintaining its structure and antigenicity. Immunogenicity of the designed nanoparticles is assessed *in vitro* and *in vivo*. In addition to introducing a novel set of reagents for multivalent display of Env trimers, this work provides both guiding principles and a detailed experimental roadmap for the generation, characterization, and optimization of Env-presenting, self-assembling nanoparticle immunogens.

## Introduction

Recombinant protein immunogens hold great promise against difficult viral and microbial targets for which there are currently no viable vaccine solutions (e.g. HIV, malaria). Well-defined structure, control over the exposure of different epitopes, high sample homogeneity and efficient manufacturing are some of the advantages of this vaccine platform (1). Engineered ectodomains of the Env glycoprotein (Env) are at the core of most present HIV vaccine development efforts (2–12). Recombinant native-like trimers, based on different HIV strains and carrying well-defined sets of stabilizing mutations, have been shown to elicit HIV-specific neutralizing antibody (NAb) responses in relevant animal models (13–16). Several of these constructs are being evaluated in human clinical trials, with many others in pre-clinical testing stages (17, 18) (ClinicalTrials.gov Identifiers: NCT03961438, NCT03816137, NCT03699241, NCT04046978).

While recombinant, native-like trimers represent a breakthrough in HIV vaccine development, identifying the most appropriate way to present them in order to maximize immunogenicity in different formulations is a major challenge that remains to be addressed. Interactions with elements of the innate and adaptive immune system are highly dependent on pathogen/immunogen shape and size, as well the distribution of surface antigens (1, 19, 20). Multivalent, particulate antigen presentation is important for several reasons. (1) A regular array of appropriately spaced antigens can lead to strong, avidity-enhanced interactions with B-cell receptors (BCRs), resulting in more robust activation of antigen-specific B cells (21–25). This factor is of particular importance for initial recruitment of immunologically naïve B cells with low affinity towards the antigen. (2) Multivalent presentation of glycosylated antigens such as HIV Env leads to more efficient crosslinking and opsonization by mannose-binding lectin (MBL), triggering the lectin pathway of the complement system (26, 27). The recruitment of complement components facilitates recognition by antigen presenting cells (APC), enhancing uptake by dendritic cells (DC) and macrophages at the site of injection and by lymph node-resident DCs, resulting in better priming of effector T cells (28). (3) Furthermore, antigen coating by MBL and complement components leads to improved trafficking through the layer of subcapsular sinus macrophages coating the lymph node, and greater accumulation of antigen within the lymph node follicles (26, 27, 29, 30). (4) Finally, particulate antigens with diameters in the 40-100 nm range require more time to penetrate the extracellular matrix around the site of injection to reach the lymphatic system, which results in slower release and hence prolonged antigen exposure (25, 28, 29, 31, 32). Extended exposure to HIV Env antigens when osmotic pumps were used for controlled immunogen release was recently correlated with more robust B-cell responses, enhanced somatic hypermutation and a greater diversity of the elicited polyclonal antibody response (33).

Two-component, self-assembling nanoparticles represent a versatile platform for presentation of HIV Env and other viral glycoproteins in a precisely defined manner (34–38). A combination of symmetric protein modeling and RosettaDesign (39–42) is applied to generate particles of appropriate geometry consisting of two oligomeric building blocks, at least one of which is genetically fused to the antigen (antigen-bearing component). The second component is generally based on a different oligomeric protein scaffold and is essential for assembly but it is typically not used for antigen presentation (assembly component). The two nanoparticle components can be purified independently, with assembly occurring when they are combined at an appropriate stoichiometric ratio. This strategy enables a high level of control over the structural and antigenic integrity of specific epitopes on the Env antigen (37, 38). In addition to simply achieving multivalent display, the geometry and spacing of presented antigens could also be varied through design (35, 36, 38). This may be particularly important for vaccine design efforts focusing on a specific set of HIV Env epitopes, which need to be presented in an appropriate and accessible orientation.

Alternative methods for the multivalent presentation of HIV Env trimers include virus-like particles (43–45), liposomes (46–49); synthetic nanoparticles (50–53); and one-component nanoparticles based on natural protein scaffolds such as lumazine synthase (54, 55), dihydrolipoylacetyltransferase (56) and ferritin (56–58). These various, complementary approaches provide additional options for immunogen design with respect to geometry and antigen distribution.

Recently, a library of two-component nanoparticles of different symmetries (tetrahedral, octahedral and icosahedral) was designed *de novo* to support the display of viral glycoprotein antigens, and found to stably present RSV F, HIV Env, and influenza HA trimers. (38). Here, we perform a detailed characterization of the designed HIV Env nanoparticle immunogens, based on BG505 and consensus group M (ConM) Env constructs. The capacity to appropriately present BG505-SOSIP trimers (5, 6) was assessed by testing the expression, assembly, structural and antigenic properties of the resulting nanoparticles and presented antigens. We then used rabbits to evaluate the immunogenicity of a particularly promising tetrahedral nanoparticle construct (T33_dn2) displaying the ConM-SOSIP trimer, in comparison to the same trimers delivered as soluble proteins.

## Results

### Nanoparticle Library

A library consisting of five self-assembling nanoparticle candidates was designed for display of SOSIP-based trimeric ectodomains of HIV Env (38). The library includes three tetrahedral (T33_dn2, T33_dn5 and T33_dn10), one octahedral (O43_dn18) and one icosahedral (I53_dn5) system (Supplementary Figure S1.a). The tetrahedral nanoparticles present 4 Env trimers when Env is fused to one of the nanoparticle components while their octahedral and icosahedral counterparts carry 8 and 20, respectively (Supplementary Figure S1.b). In each case, the antigen-bearing component (shown in orange in Supplementary Figure S1.a) was based on one of two trimeric, helical repeat protein scaffolds – 1na0C3_2 (38) or 1na0C3_3 (59) – and its N-terminus was genetically fused to the C-terminus of the Env glycoprotein ectodomain. However, the amino acid sequence of each antigen-bearing component differs between the individual constructs (Supplement Table I), with the principal differences being in the residues comprising the computationally designed interface that drives assembly in the presence of the assembly component.

### Antigen-bearing component production and characterization

We selected a BG505-SOSIP trimer as the HIV Env model for initial optimization steps. The trimer was modified by incorporating the SOSIP.v5.2 and MD39 stabilizing mutations (3, 5). A further modification involved knocking in glycosylation sites at positions N241 and N289, to occlude the immunodominant glycan hole that is present in the BG505 trimer but generally rare in other HIV strains (16, 60). Together, these sequence changes raise the thermal stability of the trimer, increase its expression levels and may improve its ability to induce more broadly active NAbs by suppressing narrow-specificity responses (5, 61). This construct is hereafter referred to as BG505-SOSIP.v5.2(7S). Trimeric building blocks of the five nanoparticle candidates were genetically fused to the C-terminus of BG505-SOSIP.v5.2(7S) to generate the various antigen-bearing components, which were expressed as secreted proteins in 293F cells. Compared to the unmodified BG505-SOSIP.v5.2(7S), all five of the engineered constructs were less efficiently expressed (Table I and Supplementary Figure S2.a). However, the four constructs that could be expressed and purified (see below) and the parental BG505-SOSIP.v5.2(7S) trimer all had very similar melting temperatures suggesting that C-terminal fusion of 1na0C3_2/1na0C3_3-based constructs does not destabilize the SOSIP antigen (Table I).

**Table 1:** Properties of BG505-SOSIP-fused nanoparticle components

| Construct | Protein yield after SEC<br>(mg per 1L of 293F cells) | T<sub>m</sub><br>(°C) | gp120 – gp41<br>cleavage | Nanoparticle<br>assembly |
| --- | --- | --- | --- | --- |
| BG505-SOSIP.v5.2(7S) | 1.5 ± 0.5 | 76.9 ± 0.5 | Full | - |
| BG505-SOSIP-T33_dn2A | 0.7 ± 0.2 | 77.9 ± 0.5 | Full | Yes |
| BG505-SOSIP-T33_dn5A | 1.0 ± 0.2 | 77.2 ± 0.5 | Full | No |
| BG505-SOSIP-T33_dn10A | 0.3 ± 0.1 | 77.5 ± 0.5 | Full | Yes |
| BG505-SOSIP-O43_dn18B | 0 | N/A | N/A | N/A |
| BG505-SOSIP-I53_dnB | 0.4 ± 0.1 | 77.7 ± 0.5 | Full | Yes |

Among the antigen-bearing components evaluated, BG505-SOSIP-O43_dn18B had a high propensity towards self-aggregation, and no trimer could be recovered after purification by size-exclusion chromatography (SEC; Supplementary Figure S2.a). Although BG505-SOSIP-T33_dn5A expressed quite efficiently, it failed to co-assemble into a nanoparticle when mixed with the corresponding assembly component, T33_dn5B (Table I, Supplementary Figure S2.c). These constructs were therefore not pursued further. BG505-SOSIP-T33_dn2A, -T33_dn10A and -I53_dn5B were expressed at ∼20-50% of the BG505-SOSIP.v5.2(7S) trimer yield and efficiently assembled into nanoparticles (Table I, Supplementary Figure S2.c). The yield reductions were probably due to self-aggregation, as indicated by SEC profiles (Supplementary Figure S2.a). The absence of appropriate assembly component during expression leads to solvent exposure of nanoparticle interface residues in the antigen-bearing component that are mainly hydrophobic, which is the most probable cause of the observed self-aggregation. The above three candidates were then further characterized. An SDS-PAGE analysis showed that the SEC-purified samples were homogeneous and cleaved into gp120 and gp41 (+ nanoparticle component) subunits that were separated when a reducing agent was added to break the engineered disulfide bonds (Figure 1.a). Analysis of 2D class-averages from negative stain electron microscopy (NS-EM) imaging suggests that the BG505-SOSIP trimer components are in a native-like conformation, with the density corresponding to the fused nanoparticle component clearly discernible (Figure 1.b; see red arrows). A low level (∼ 8 %) of monomeric protein was present only in the BG505-SOSIP-T33_dn10A sample.

**Figure 1.**
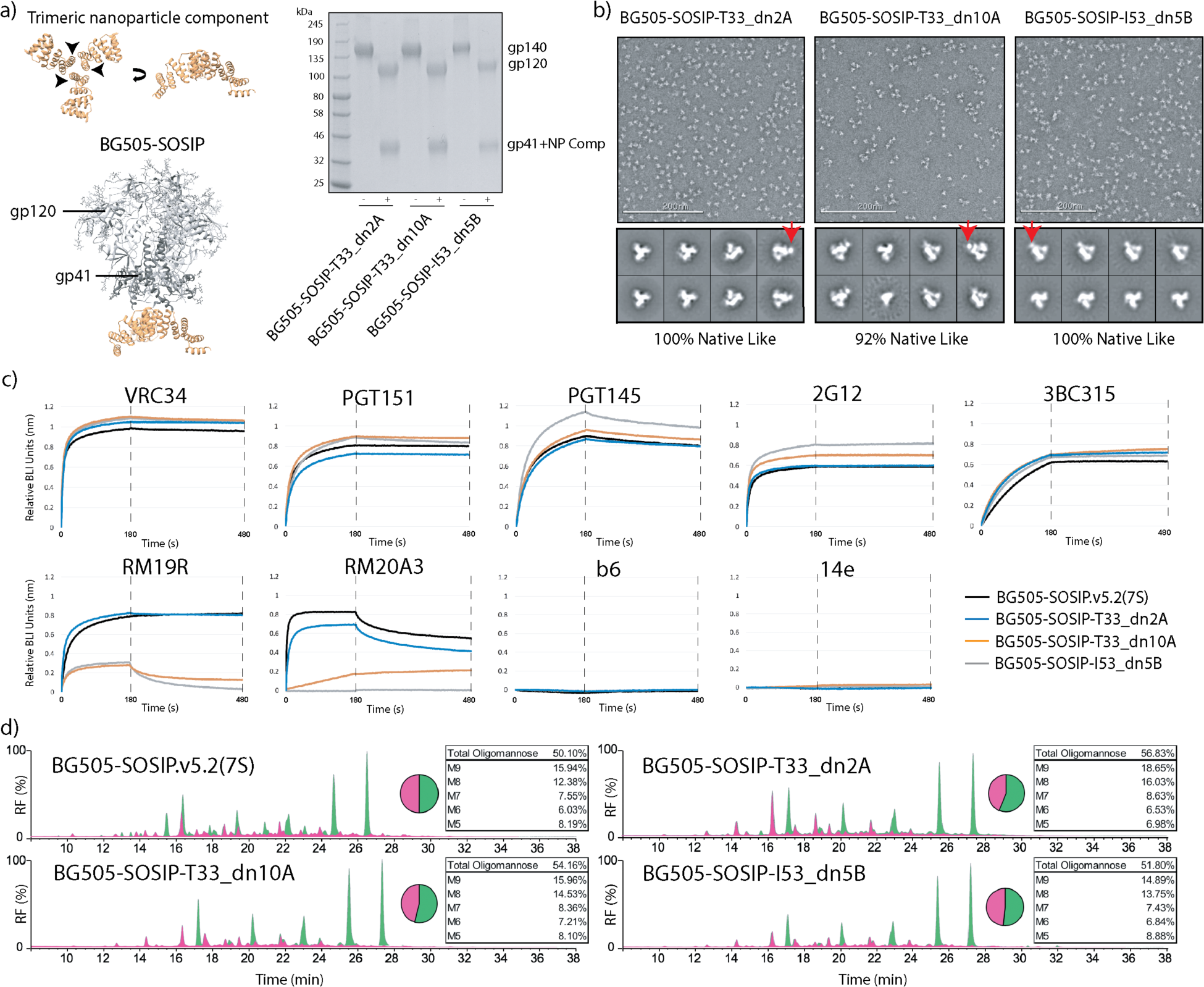
Evaluation of the antigen-presenting components. (a) Antigen-bearing components were generated by fusing the N-termini of trimeric nanoparticle building blocks to BG505-SOSIP.v5.2(7S) (*left*). The SDS-PAGE gel of the purified antigen-bearing components denatured in the presence (+) and absence (-) of reducing agent (*right*). (b) NS-EM analysis of the purified antigen-bearing components (representative raw micrograph and 2D class averages). Red arrows indicate the location of the fused nanoparticle component. Analysis of the resulting 2D classes suggests that the Env antigen is in a native-like, trimeric form in all three antigen-bearing components. A small percentage of monomer/dimer-like particles were detected in the BG505-SOSIP-T33_dn10A sample (2^nd^ class from the left in the bottom row). (c) BLI analysis of the antigenicity of three antigen-bearing components compared to BG505-SOSIP.v5.2(7S). (d) Glycan composition analysis for the three antigen-bearing components and BG505-SOSIP.v5.2(7S). Peaks sensitive to endoglycosidase H digestion correspond to oligomannose-type glycans and are colored green.

We used Biolayer Interferometry (BLI) and a panel of antibodies (IgG) specific to different Env epitopes and conformations to probe the antigenicity of the antigen-bearing components in their trimeric forms, prior to nanoparticle assembly (Figure 1.c). All the samples interacted with neutralizing antibodies (NAbs) specific to the closed, prefusion state of the BG505-SOSIP trimer (VRC34, PGT145, PGT151, 2G12, 3BC315), but did not bind to b6 and 14e, two non-neutralizing antibodies (non-NAbs) that recognize more open trimer conformations and free monomers. The RM19R and RM20A3 antibodies showed markedly lower binding to the BG505-SOSIP-T33_dn10A and -I53_dn5B components compared to the BG505-SOSIP.v5.2(7S) trimer. These two non-NAbs interact with the base of the BG505-SOSIP.v5.2(7S) trimer, which is the site of fusion to nanoparticle component. The occlusion of these epitopes on the antigen-bearing fusion constructs is expected and potentially useful, as the induction of non-NAbs against the trimer base could be immune-distractive. Surprisingly, BG505-SOSIP-T33_dn2A displayed no change in RM19R binding compared to free BG505-SOSIP.v5.2(7S), and only a marginal decrease in binding to RM20A3.

In a glycan composition analysis, the ratio of oligomannose and complex glycans was very similar for BG505-SOSIP-T33_dn2A, -T33_dn10A, -I53_dn5B and the parent BG505-SOSIP.v5.2(7S) trimer, with the oligomannose proportion ranging from 50% to 57% (Figure 1.d). The ratio of individual oligomannose species remains similar in all samples with the highest difference being present in BG505-SOSIP-T33_dn2A, where the relative content of oligomannose species of higher molecular weight (M8 and M9) is slightly elevated.

A site-specific glycan composition analysis showed that glycan processing was conserved at key epitopes on all the trimer samples, with sites such as N332 and N160 containing predominantly oligomannose-type glycans (Supplementary Figure S3.). One major difference was, however, visible at gp41-site N637 on BG505-SOSIP-T33_dn2A, which contains 100% oligomannose-type glycans compared to 34% on BG505-SOSIP.v5.2(7S). Additionally, this site is fully glycosylated in the three antigen-bearing components, and only partially glycosylated (56%) on the BG505-SOSIP.v5.2(7S) trimer. Glycan sites on gp120 located at the protomer interfaces of the BG505-SOSIP-T33_dn2A and -T33_dn10A components were also enriched for oligomannose-type glycans, most notably N276 and N355. Specifically, the oligomannose contents of 67% for N276 and 22% for N355 on BG505-SOSIP.v5.2(7S) increased to >95% for N276 and >75% for N355 on the BG505-SOSIP-T33_dn2A and -T33_dn10A components. The V1/V2 glycan sites N133, N137 and N185e were <95% occupied on all the trimer constructs, while the N156 site on BG505-SOSIP.v5.2(7S) was 89% occupied. Among the closely spaced glycosylation sites at N289, N295 and N301, N289 and to a lesser extent N301 glycosylation appear to be affected by the fusion to the antigen-bearing component of the nanoparticle. A reduction of 13-19% was observed in the occupancy at N289 site in the three tested antigen-bearing components when compared to the parental BG505-SOSIP.v5.2(7S).

Collectively, these data indicate that several of the de novo-designed nanoparticle trimers were able to present a genetically fused native-like Env trimer without major changes to antigen structure, stability, or glycan profile.

### Nanoparticle assembly, antigenic and structural characterization

The three antigen-bearing components described above (BG505-SOSIP-T33_dn2A, - T33_dn10A and -I53_dn5B) were then tested for nanoparticle assembly (Figure 2.a). The corresponding assembly components, T33_dn2B, T33_dn10B and I53_dn5A required for nanoparticle formation were expressed in *E. coli* and purified as described in the Methods section. Analysis of the purified assembly components by SDS-PAGE is shown in Supplementary Figure S2.b. Equimolar amounts of the antigen-bearing component and the corresponding assembly component (on a subunit:subunit basis) were combined and incubated for 24 hours at three different temperatures (4, 25 and 37 °C) before assembly was evaluated using native PAGE. In each case, assembly efficiency increased with the incubation temperature. The tetrahedral nanoparticles, T33_dn2 and T33_dn10, assembled at a high yield (∼80-100%) under all the conditions tested. However, the icosahedral nanoparticle, I53_dn5, assembled less efficiently. After a 24-hour incubation at 37 °C, only ∼30% of the input material migrated as nanoparticles on a native gel, though when the incubation period was extended to 72 h at 37 °C, the yield increased to ∼70% (Supplementary Figure S2.c).

**Figure 2.**
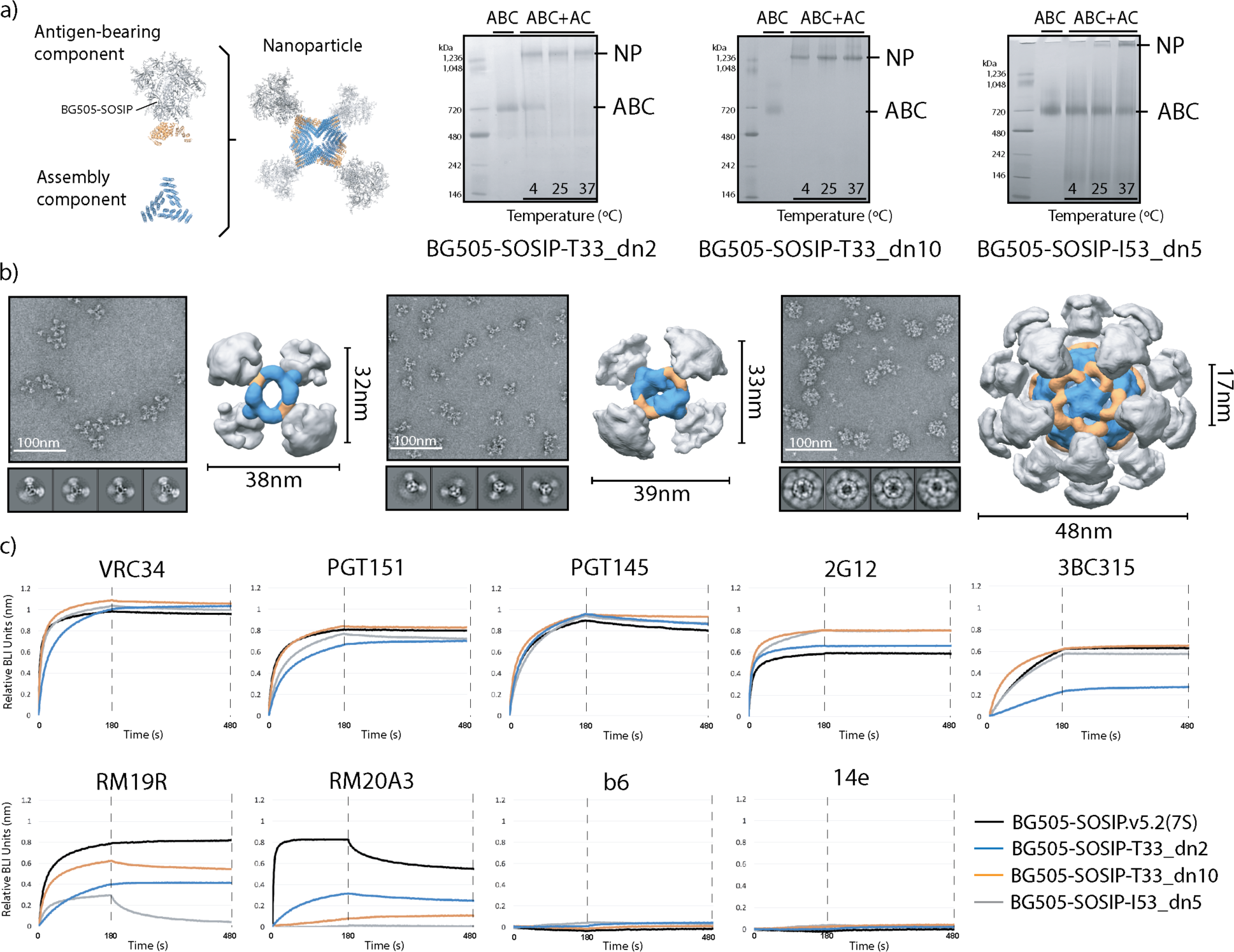
Nanoparticle assembly and characterization. (a) Schematic representation of individual components and assembled nanoparticle (*left*). Nanoparticle assembly tests were performed at different temperatures and the assembly efficacy was assessed using Native PAGE. NP, nanoparticle; ABC, antigen-bearing component; AC, assembly component; (*right*). (b) Negative stain EM analysis of purified nanoparticles. Representative raw micrographs, 2D class averages and 3D reconstructions are shown for BG505-SOSIP-T33_dn2 (left), -T33_dn10 (middle) and -I53_dn5 (right) nanoparticles. 3D maps are segmented and color-coded (BG505 SOSIP, gray; antigen-bearing component, orange; assembly component, blue). Particle diameters and the average apex-apex distance between the two closest neighboring Env trimers are shown for each nanoparticle. These data are also described in Ueda et al., Submitted. (c) BLI analysis of antigenicity of assembled nanoparticles compared to the BG505-SOSIP.v5.2(7S) trimer.

The presence of both the antigen-bearing component and the assembly component in SEC-purified nanoparticles was verified using SDS-PAGE (Supplementary Figure S2.d). Sample homogeneity and structural integrity were assessed using NS-EM. Representative raw micrographs, 2D class-averages and reconstructed 3D models (adapted from (38)) show that the assembled forms of the nanoparticles were consistent with the predictions of the computational design models, with individual building blocks clearly discernible in each case (Figure 2.b).

Nanoparticle stability under various conditions was evaluated using a native PAGE assay (Supplementary Figure S4.). The tetrahedral particles (T33_dn2 and T33_dn10) were highly stable in buffers with pH values in the range 5–9 and at NaCl concentrations from 25–1000 mM. They also remained assembled at temperatures up to 65 °C and withstood multiple freeze-thaw cycles. In contrast, the icosahedral nanoparticles were less stable under the various test conditions and were particularly sensitive to the freeze-thaw procedure. The latter observations are consistent with the presence of significant amounts of unassembled components revealed by NS-EM (Figure 2.b, right), and also with the greater difficulties in assembling the icosahedral particles (see above).

We used BLI to assess whether nanoparticle presentation interfered with the accessibility of antibody epitopes, compared to the parental BG505-SOSIP.v5.2(7S) trimer (Figure 2.c). Overall, the antigenic profiles of the assembled nanoparticles, the corresponding antigen-bearing components and the free trimer were very similar (Figure 2.c, and see also Figure 1.c). Compared to the BG505-SOSIP.v5.2(7S), the greatest decrease in binding was seen with the base-specific antibodies, RM19R and RM20A3, suggesting that nanoparticle assembly further decreases the accessibility of the trimer base (Figure 2.c). There was also a decrease in the binding of the BG505-SOSIP-T33_dn2 nanoparticles to the 3BC315 broadly neutralizing antibody (bNAb), compared to the free trimer (Figure 2.c). As no such difference was seen with the BG505-SOSIP-T33_dn2A antigen-bearing component (Figure 1.c), assembly of this nanoparticle appears to reduce the accessibility of epitopes for bNAbs such as 3BC315 that are located near the bottom of the trimer.

### Cryo-EM analysis of assembled nanoparticles

We used cryo-electron microscopy (cryo-EM) to obtain more detailed information on the structures of the BG505-SOSIP-T33_dn10 and -I53_dn5 nanoparticles and the Env trimers they display (Figure 3.). The data were processed as described in the Methods section and summarized in Supplementary Figures S5 and S6, with data acquisition and processing statistics shown in Supplementary Table II. Due to the flexible nature of the linker between the nanoparticle component and the BG505-SOSIP trimer, the initial 3D reconstructions of the complete nanoparticles generated only highly diffuse and poorly defined density for BG505-SOSIP (Figure 3. and Supplementary Figure S5. and S6.). The data were therefore computationally segmented to reconstruct the nanoparticle core and the displayed trimer as two independent but flexibly linked entities (sub-particles).

**Figure 3.**
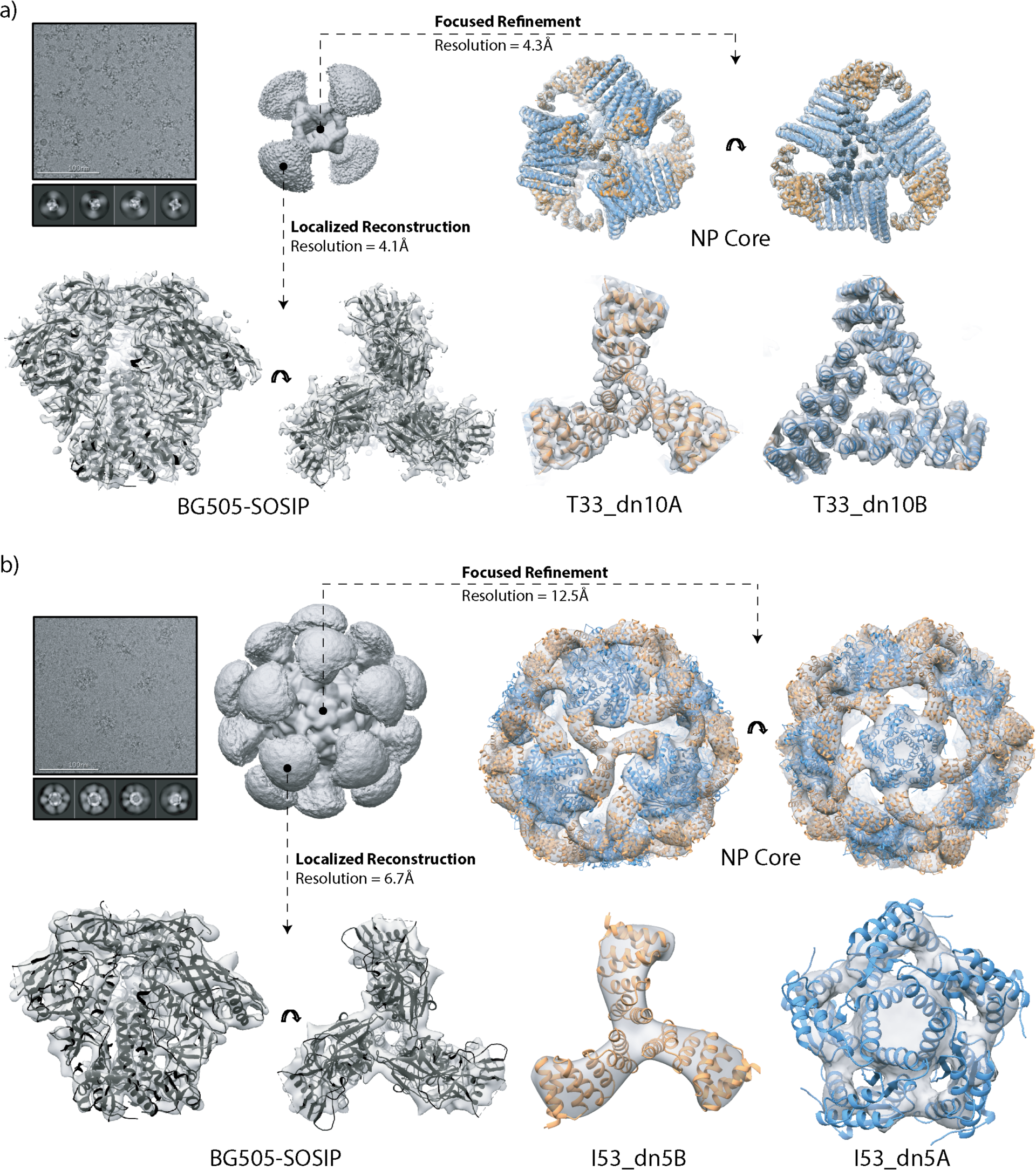
Cryo-EM analysis of tetrahedral and icosahedral nanoparticles. (a) Tetrahedral BG505-SOSIP-T33_dn10 nanoparticle; (b) Icosahedral BG505-SOSIP-I53_dn5 nanoparticle. Sample micrograph, 2D class averages and initial 3D reconstructions of the full nanoparticles are displayed in the top left part of the corresponding panels. Focused refinement was applied to generate a 3D reconstruction of the nanoparticle core (top and bottom right, maps are in light gray). The refined model of T33_dn10 and the Rosetta_design model of I53_dn5 are docked into the corresponding maps (antigen-bearing component, orange; assembly component, blue). Localized reconstruction approach was used for analysis of the presented antigen (bottom left). Refined BG505-SOSIP models are shown in black.

Nanoparticle reconstruction was performed using a focused refinement procedure in which a solvent mask around the nanoparticle core excluded the signal originating from BG505-SOSIP trimers. Symmetry was applied in all 3D classification and refinement steps: tetrahedral for T33_dn10 and icosahedral for I53_dn5. The final resolutions of the reconstructed maps were 4.25 Å and 12.50 Å for T33_dn10 and I53_dn5, respectively. The T33_dn10 core design model from Rosetta Design was relaxed into the EM map using a combination of Rosetta relaxed refinement (62) and manual refinement in Coot (63). Model refinement statistics are shown in Supplementary Table III. The model-to-map fit for the refined structure is shown in Figure 3.a (right). There are only small, local structural differences between the Rosetta-predicted, unliganded (38) and BG505-SOSIP-bearing models of the T33_dn10 -nanoparticle core. The C*α* RMSD between the experimentally determined and predicted models is 1.43 Å (on the level of the asymmetric unit), demonstrating that the presence of four displayed BG505-SOSIP trimers did not cause any major structural rearrangements within the nanoparticle core. Model refinement was not performed for the I53_dn5 core because the resolution was too poor. The design model of I53_dn5 was, however, highly concordant with the reconstructed EM map (Figure 3.b, right).

The nanoparticle-displayed BG505-SOSIP trimers were reconstructed using a localized reconstruction approach (64). The signal corresponding to the nanoparticle core was subtracted from the particle images and the trimer sub-particles were extracted and aligned independently. The final BG505-SOSIP trimer subset from the T33_dn10 nanoparticle dataset consisted of 84,435 sub-particles and was reconstructed to 4.14 Å resolution. A model of the displayed BG505-SOSIP was refined using a combination of Rosetta-relaxed and manual refinement in Coot (Figure 3.a, Supplementary Table III). The close agreement of the refined model with published structures of the BG505-SOSIP trimer (PDB entries: 5CEZ and 5ACO (65, 66)) implies that nanoparticle assembly does not interfere with the structural integrity of the trimer. For the I53_dn5 nanoparticle dataset, the final subset included 7,737 sub-particles and was reconstructed to a resolution of 6.67 Å. The quality of the BG505-SOSIP map was significantly better than the corresponding I53_dn5 nanoparticle core but still too poor resolution to refine a structural model. However, a published structure (PDB entry: 5CEZ (65)) was docked into the map and exhibited a good fit (Figure 3.b).

### Nanoparticle presentation affects trimer interaction with B cells and immunogenicity

BG505-SOSIP-T33_dn2, -T33_dn10, and -I53_dn5 nanoparticles were then functionally analyzed as immunogens. Their capacity to stimulate antigen-specific B cells was evaluated using K46 mouse B-cell lines that expressed IgM versions of three different HIV-specific bNAbs on their surfaces (PGT145, VRC01 and PGT121; Figure 4.). The B cells were treated with equimolar amounts of BG505-SOSIP.v5.2(7S) antigen presented as free trimers, antigen-bearing components or assembled nanoparticles, and the relative number of cells responding to each antigen was quantified (Figure 4.). Ca^2+^ mobilization inside B cells, measured by fluorescence-activated cell sorting (FACS), was used as an indicator of antigen-induced activation. For clarity, the data from each B cell line are presented in two panels, the first showing the different antigen-bearing trimeric components, and the second the assembled nanoparticles. The free trimers (unmodified BG505-SOSIP.v5.2(7S) and the antigen-bearing components) induced only very low levels of Ca^2+^ flux (Figure 4., top) suggesting that the local antigen concentration was insufficient for efficient BCR crosslinking and B-cell activation (67). In contrast, presenting the same trimers on the surface of each of the three nanoparticles activated the B cells much more strongly, as quantified by the higher percentage of cells in which a Ca^2+^ flux occurred (Figure 4., bottom). The icosahedral BG505-SOSIP-I53_dn5 nanoparticle triggered stronger Ca^2+^ signal than the two tetrahedral nanoparticles in B cells expressing PGT145. In VRC01-expressing cells this difference was less pronounced, particularly when compared to BG505-SOSIP-T33_dn10. Although the signal-to-noise ratio was generally low in PGT121-expressing cells, there was an increased Ca^2+^ flux response with the nanoparticles compared to free trimers.

**Figure 4.**
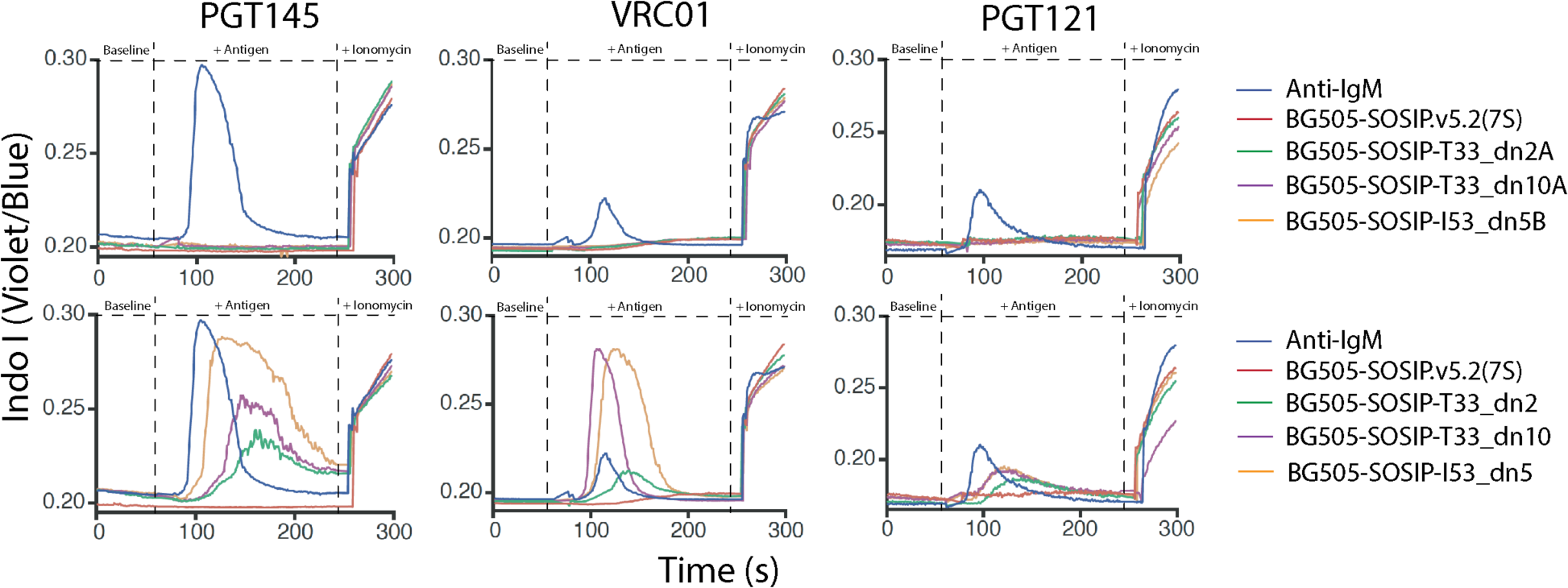
B-cell activation by trimeric components and nanoparticles. Ca^2+^ flux (Indo I fluorescence) was used to assess the activation of B cells expressing HIV Env-specific IgM receptors (PGT145, VRC01 and PGT121) by equimolar amounts of BG505-SOSIP in the form of free trimers, fused to the antigen-bearing components (top row) and on the surface of assembled nanoparticles (bottom row). The antigen was introduced 60 s after the start of each measurement. Ionomycin was added after 240 s. Anti-IgM antibody was used as a positive control.

Based on the totality of the production, antigenicity and biophysical data, we determined the tetrahedral T33_dn2 nanoparticle to be the best system for displaying the BG505-SOSIP trimer. We then assessed whether this nanoparticle design could also be used to present SOSIP trimers based on another HIV Env sequence, specifically consensus group M (ConM) (8). The ConM-SOSIP trimers are being evaluated clinically (ClinicalTrials.gov Identifiers: NCT03961438, NCT03816137) and have been studied in the context of the designed, two-component nanoparticle, I53-50 (37). The ConM-SOSIP construct used in the experiments, termed ConM-SOSIP.v7, was engineered to include the SOSIP.v5.2 and TD8 stabilizing mutations (see the Method section and Supplementary Table I for sequence information) (8, 37, 68). The trimeric ConM-SOSIP-T33_dn2A component expressed at a similar level as its BG505-based counterpart (∼ 0.6 mg per 1 L of 293F cells after SEC, Supplementary Figure S7.a), and was assembled efficiently into nanoparticles when incubated with equimolar amounts of the T33_dn2B component at 4 °C (> 90% assembly after 24 h, Supplementary Figure S7.b). The purified ConM-SOSIP-presenting nanoparticles were characterized by NS-EM and Surface Plasmon Resonance (SPR) (Supplementary Figure S7.b and c). The NS-EM analysis showed that the ConM-SOSIP-T33_dn2 nanoparticles assembled to the target tetrahedral architecture and were highly homogeneous. Assessed by SPR, the ConM-SOSIP.v7 trimers on their surfaces retained the capacity to interact with trimer-specific antibodies. An antigenicity comparison with equimolar amounts of the free ConM-SOSIP.v7 trimer suggested that the nanoparticles efficiently presented NAb epitopes located on the upper half of the trimer but that the accessibility of epitopes located towards the bottom of the trimer (i.e. the fusion peptide, 35O22 and 3BC315 epitopes) was partially impaired. This data is consistent with the similar SPR-based epitope accessibility studies performed on T33_dn2 nanoparticles presenting BG505-SOSIP.v5.2(7S) (38).

The ConM-SOSIP-T33_dn2 nanoparticles were then tested for immunogenicity in New Zealand White Rabbits and compared to the soluble ConM-SOSIP.v7 trimer (Figure 5.). Two groups of 5 rabbits were used, with the immunogen dose adjusted to ensure that all the animals received an equimolar amount of ConM-SOSIP.v7 trimer (30 µg). The rabbits were immunized at weeks 0, 4 and 20, with blood draws performed at weeks 0, 2, 4, 6, 8, 12, 16, 20 and 22. Immune responses were monitored in sera by measuring ConM-SOSIP trimer-specific antibody titers, by ELISA, and NAb titers using the TZM-bl cell assay.

**Figure 5.**
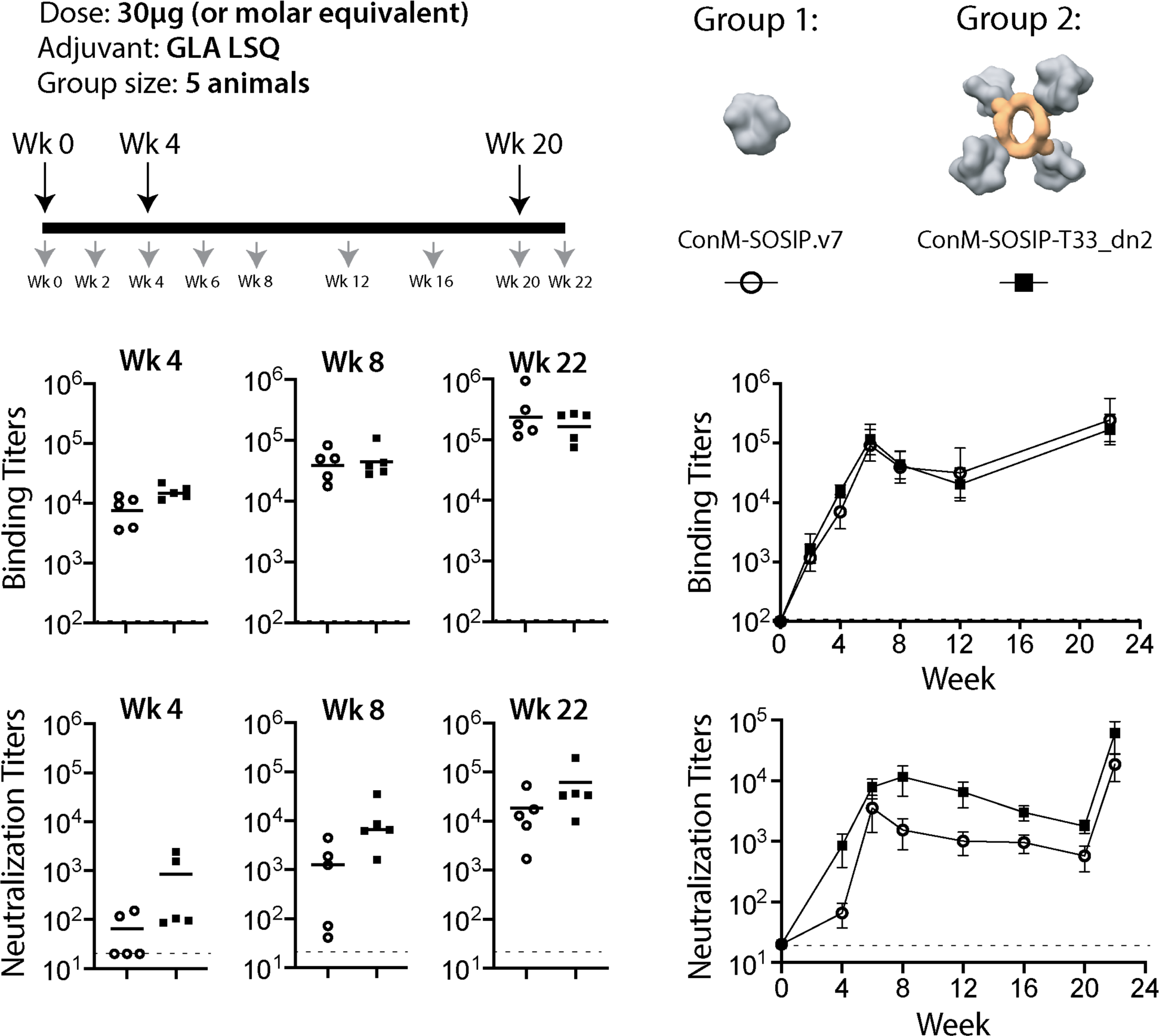
Immunogenicity of nanoparticle-presented ConM-SOSIP. Two groups of 5 rabbits were immunized with 30 µg of soluble ConM-SOSIP.v7 trimer (Group 1, open circles) or the equivalent amount presented on the T33_dn2 tetrahedral nanoparticle (Group 2, black squares). The immunization (large arrows) and bleed (small arrows) schedules are shown on the top left, and depictions of the immunogens on the top right. The serum anti-trimer binding and neutralizing antibody titers in each animal at weeks 4, 8 and 22 are presented as scatter plots with the mean titers indicated by lines. The mean titers for each group are plotted longitudinally on the right with the bars corresponding to the standard error of the mean. An AUC statistical analysis (based on two-tailed Mann-Whitney U-test) of titer values in Group 1 versus Group 2 as a function of time results in p values of 0.69 and 0.056 for the binding and neutralization titers, respectively. The data for Group 1 (ConM-SOSIP.v7) were adapted from Brouwer et al., 2019 (see Methods section for details).

Both immunogens induced anti-ConM-SOSIP binding antibodies (Figure 5. middle panels). The mean ELISA binding titers were comparable between the immunization groups at each time point. An area under the curve (AUC) statistical analysis using a two-tailed Mann-Whitney U-test generated a p-value of 0.69, implying that there is no statistically significant difference in the binding titers between the two groups across all time points. Antibody responses of similar magnitude (week-22 ELISA titers of ∼10^5^) against the nanoparticle core were also detected in the ConM-SOSIP-T33_dn2 immunogen group but not the soluble trimer group (Supplementary Figure S8.a).

NAb titers against the autologous Tier-1 ConM virus were more variable than the binding antibody responses, both within and between the immunization groups (Figure 5, bottom panels). Within each group, the NAb titers spanned a range of ∼100-fold at each time point, a greater variation than the ∼10-fold range in the anti-trimer titers. Every rabbit immunized with ConM-SOSIP-T33_dn2 nanoparticles generated a detectable NAb response (titer >20) after the first immunization, and the mean titers were consistently higher in this group than in the ConM-SOSIP.v7 group (p = 0.016 by AUC analysis) (Supplementary Figure S8.b). Taken together, the serological data suggest that presenting the ConM-SOSIP.v7 trimer on the T33_dn2 nanoparticle increased the proportion of the B-cell response directed against the autologous NAb epitope(s).

We next employed EM-based polyclonal epitope mapping (EMPEM) (69) to characterize the specificities of antibodies elicited by the two immunogens (Figure 6). Polyclonal Fab samples were prepared from week-22 sera from the two animals in each group with the highest ConM NAb titers (Group 1, r2381 and r2382; Group 2, r2383 and r2385), and also from the same animals at week 4. The purified Fabs were complexed with ConM-SOSIP.v9 trimers, which contain additional stabilizing mutations (see Methods and Supplementary Figure S9. a and b). Raw data, sample micrographs, 2D class averages and reconstructed 3D maps of the trimer-Fab complexes are shown in Supplementary Figure S10. Composite figures based on these datasets are shown in Figure 6, where polyclonal Fabs of the various specificities detected by the 3D analysis are docked onto a reference SOSIP trimer model. Fab epitopes are defined based on the partial overlap with the receptor binding site (CD4bs), variable loops (e.g., V1, V2, V3 and their combinations) and, in some cases, overlap with one or more glycan sites (e.g., N611, N618/N625 and N355/N289). For reference, binding and neutralizing antibody titers are also shown for each of the four animals at the two time-points.

**Figure 6.**
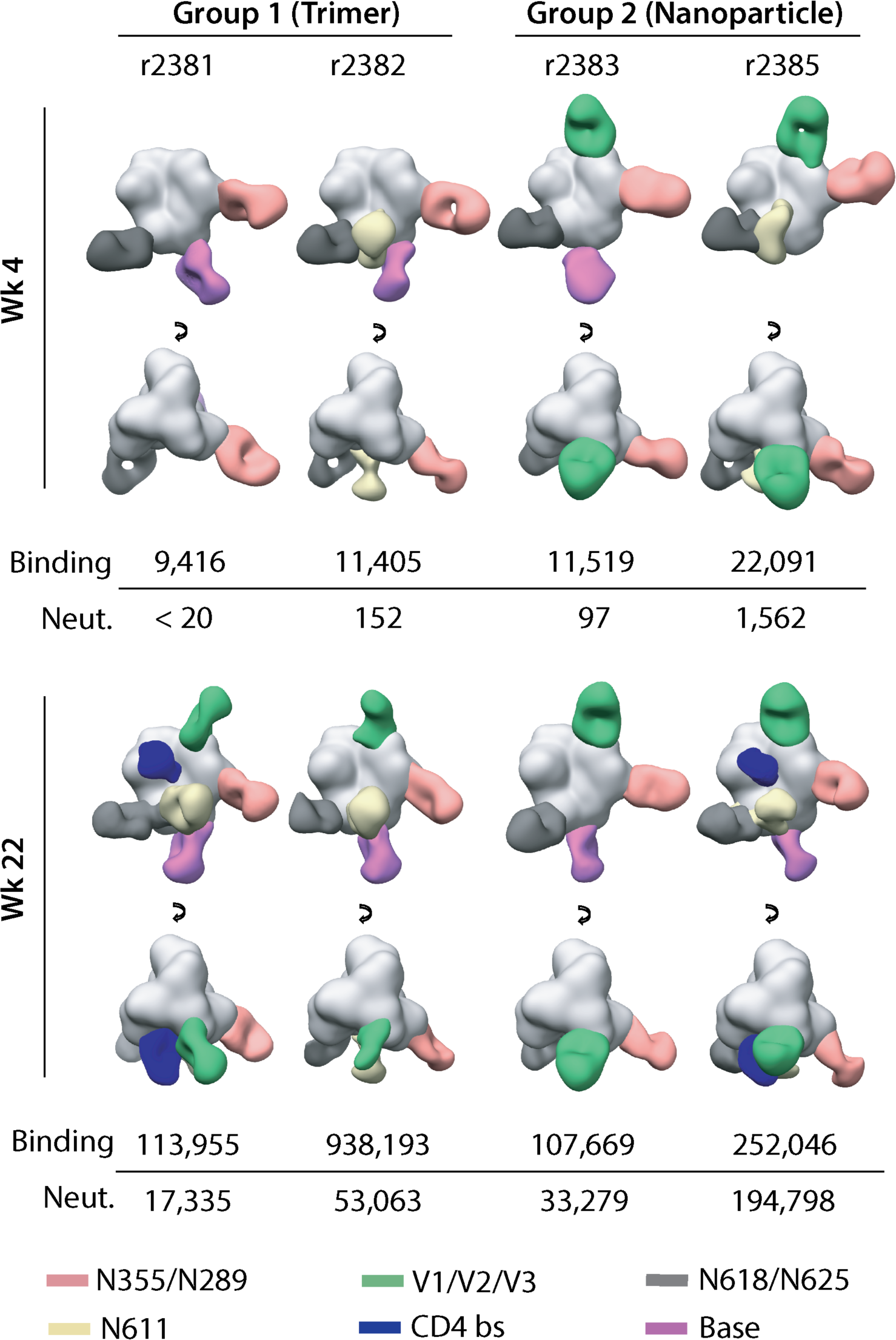
EMPEM analysis of antibody responses in immunized animals. Composite figures generated from EMPEM analysis performed using sera collected from the two animals in each group that have the highest ConM NAb titers at week 22. Data are shown for the week 4 (post-prime) and week 22 (post-final boost) time points. For simplicity, only a single antibody is shown for each epitope cluster. Epitope definitions are summarized in the text and color coded as indicated at the foot of the figure. The anti-trimer binding antibody and neutralizing antibody titers for each serum sample are listed below the images.

On a qualitative level, the data indicate that the immunogenic properties of the ConM-SOSIP.v7 trimer were not influenced by how it was presented. This inference is particularly true at week 22 (2 weeks after the final boost) where similar antibody specificities were detected in all 4 four rabbits, notably ones targeting the V1/V2/V3 interface, the N618/N625 and N355/N289 glycan epitopes and the trimer base. The N611 glycan epitope was also targeted in three animals (r2383 was the exception), while CD4bs-associated responses were detected in one animal from each group (r2381 and r2385). The CD4bs is a well-characterized bNAb target that is the focus of several vaccine design strategies (54, 70). We were unable to identify any relationship between the type and quantity of the antibody specificities induced and the binding and NAb responses present in the corresponding sera at the week-22 time point.

There was more variability in the antibody specificities detected at week 4. The N618/N625 and N355/N289 glycan epitopes were targeted in all 4 animals, while anti-base antibodies were induced in both animals from the soluble trimer group and in animal r2383 from the nanoparticle group. Antibodies to the N611 glycan site were detected in one animal from each group. However, antibodies against the V1/V2/V3 interface, located at the top of the trimer, were visible only in the nanoparticle immunogen group at this early time point. The initial antibody response induced after a single immunization (i.e., at week 4) may be skewed towards the V1/V2/V3 epitopes at the trimer apex and away from the trimer base in the nanoparticle immunization group, compared to the soluble trimer group. The early differences in how the antibody responses are primed may contribute to the higher neutralization titers induced in the nanoparticle group after three immunizations (i.e., by week 22).

## Discussion

We evaluated the structure, stability, antigenicity and immunogenicity of two-component nanoparticles presenting HIV Env trimers. In contrast to previous work, the nanoparticles were designed *de novo* to comprise a trimeric building block tailored to support C-terminal fusion to HIV Env and other trimeric viral antigens (38). Our library consisted of five nanoparticle candidates of different geometries: three tetrahedral, one octahedral and one icosahedral, presenting 4, 8 and 20 Env trimers, respectively. Two tetrahedral (T33_dn2 and T33_dn10) and one icosahedral nanoparticle (I53_dn5) were able to support the presentation of BG505-SOSIP.v5.2(7S), the model Env trimer selected for optimization. Biophysical and antigenic characterization of the nanoparticle-presented trimers showed that they were folded appropriately, with no detectable adverse effects of nanoparticle incorporation. The nanoparticle-displayed trimers were also glycosylated comparably to their soluble counterparts (18, 71). However, nanoparticle presentation did reduce the accessibility of epitopes on and proximal to the trimer base, which is consistent with antibody binding analyses of other self-assembling nanoparticle designs (37, 38).

The tetrahedral nanoparticles (T33_dn2 and T33_dn10) are the lowest valency nanoparticle systems developed for presentation of Env trimer immunogens. Both assembled efficiently and were highly stable under a range of stress-inducing conditions in vitro. They were also superior to soluble trimers when tested in Ca^2+^ flux-based assays of B-cell stimulation. This finding implies that that as few as four BG505-SOSIP antigens are sufficient to meet the minimum threshold for activation when B-cell receptors have high affinity for the antigen (Figure 4.). Of note is that the BG505- and ConM-SOSIP-T33_dn2 nanoparticles can be produced at high yield; soluble trimers based on these two genotypes are currently being evaluated for safety and immunogenicity in human clinical trials (ClinicalTrials.gov Identifiers: NCT03961438, NCT03816137, NCT03699241).

The observed issues with assembly and stability of the icosahedral BG505-SOSIP-I53_dn5 nanoparticles are somewhat unexpected given that these particles assemble very efficiently when Env is not present (assembly completes within ∼30 minutes) and when other antigens (i.e., influenza HA and prefusion RSV F) are displayed by genetic fusion (38). The instability is most likely caused by the crowding of heavily glycosylated Env trimers on the surface of I53_dn5. Additionally, excessive steric clashes could lead to the formation of partially assembled particles that are difficult to distinguish and purify away from the fully assembled ones. It is possible that partially assembled particles constitute the majority of the sample following purification, and that these were unable to withstand the stresses to which they were subjected. Designing nanoparticles with larger diameters to decrease the antigen crowding effect while maintaining the appropriate spacing necessary for multivalent interactions with B cells may help achieve more optimal properties with the higher valency particles.

We used rabbits to compare the immunogenicity of the tetrahedral ConM-SOSIP T33_dn2 nanoparticles and the corresponding soluble ConM-SOSIP.v7 trimers (Figures 5 and 6). The anti-trimer binding antibody responses were similar in the two groups, but the autologous NAb titers were higher in the nanoparticle recipients. Polyclonal epitope mapping suggested that this could be due to better shielding of epitopes that are targets for non-neutralizing antibodies (e.g. the trimer base) and/or more efficient priming of antibodies targeting the variable loops located on the top of the ConM trimer (V1/V2/V3). This is supported by the earlier immunogenicity studies performed with ConM-based immunogens that identified variable loops at the ConM apex (V2 and V3) as the most dominant neutralizing epitopes (8, 37). Furthermore, a meta-analysis of multiple nanoparticle immunization experiments shows that the greatest benefit, compared to soluble trimers, arises when the NAb epitopes are located near the trimer apex (19). Our data supports this conclusion and suggests that immunization platforms based on apex-targeting Env immunogens may benefit from nanoparticle display. Importantly, however, the immunogenicity data confirms that non-apex epitopes are also accessible and that T33_dn2-presented Env trimers are capable of eliciting antibodies against the same epitopes as free trimers. This provides a rationale for using tetrahedral nanoparticles with a wide range of epitope-focused HIV vaccine design approaches.

The anti-base response seen in the nanoparticle group was unexpected; it most likely arises from partial nanoparticle disassembly *in vivo*, a supposition supported by the relatively steep angle with which these antibodies approach the trimer. More studies are required to understand the processes that induce nanoparticle disassembly, the kinetics of this process and the impact on trimer immunogenicity.

Overall, our data confirms the ability of designed two-component nanoparticles to at least moderately improve the immunogenicity of HIV Env antigens and provides *in vitro* and *in vivo* evidence of the potential benefits of these systems. Additionally, we introduce and validate two tetrahedral nanoparticle platforms that can be used as immunogens as well as tools for basic structural and immunological experiments. The screening, biophysical and structural characterization approach that we describe provides a roadmap for generating and evaluating multivalent Env trimer immunogens based on two-component self-assembling scaffolds.

## Materials and Methods

### DNA vectors and cloning

Constructs containing BG505-SOSIP.v5.2(7S) or ConM-SOSIP.v7 genes codon-optimized for mammalian cell expression were subcloned into a pPPI4 vector. BamHI and NheI restriction sites were used for insertion of different C-terminal nanoparticle assembly components. Restriction enzymes (BamHI-HF and NheI-HF) and Quick Ligation kit were purchased from New England Biolabs (NEB). Linker insertion and modifications in the antigen-bearing components were achieved using Q5 Site Directed Mutagenesis Kit (NEB). Custom DNA primers produced by Integrated DNA technologies (IDT). Assembly component DNA constructs, codon optimized for bacterial expression were subcloned into a pET28b (+) vector. Protein sequences of all constructs used in this study are shown in Supplement Table I.

### Expression and purification of BG505-SOSIP, ConM-SOSIP and antigen-bearing components

BG505-SOSIP construct used in this study was engineered to carry a combination of SOSIP.v5.2 (mutations: A501C, T605C, I559P, E64K, A73C, A316W, A561C) (5) and MD39 (mutations: M271I, A319Y, R585H, L568D, V570H, R304V, F519S) (3) stabilizing mutations and glycan knock-ins at positions 241 (mutations: P240T, S241N) and 289 (mutations: F288L, T290E, P291S). ConM-SOSIP.v7 construct had SOSIP.v5.2 and TD8 stabilizing mutations (8, 68). ConM-SOSIP.v9 construct was engineered with SOSIP.v6 (5), MD39 (3) and TD8 (68) stabilizing mutations. For sequence information on ConM-SOSIP constructs see Supplementary Table I and Supplementary Figure S9.b. pPPI4 DNA vectors carrying free SOSIP trimers or SOSIP-based antigen-bearing components were transfected into FreeStyle 293F cells using polyethyleneimine (Polysciences, Inc) as described previously(6). 6 days post transfection the cells were spun down (7,000 RPM for 1 hour at 4 °C) and the supernatant was cleared by vacuum filtration (0.45 µm filtration units, Millipore Sigma). Antigen-bearing components were purified using immuno-affinity column with immobilized PGT145 IgG (Sepharose 4B resin, GE Healthcare Life Sciences). 3 M MgCl2 + 250 mM l-arginine (pH 7.2) buffer was applied for protein elution. Eluate was collected into an equal volume of the SEC buffer (25 mM Tris + 500 mM NaCl + 250 mM l-arginine + 5 % glycerol, pH 7.4). Affinity-purified protein was concentrated and buffer exchanged to SEC buffer using Amicon ultrafiltration units, 100 kDa cutoff (Millipore Sigma). Size exclusion chromatography was used as a final purification step (HiLoad 16/600 Superdex S200 pg column). Purified proteins were stored at 4 °C.

### Assembly component expression and purification

E coli expression system was used for assembly component production (T33_dn2B, T33_dn10B and I53_dn5A). BL21-DE3 cells (NEB) were transformed with pET28b (+) vector carrying the appropriate gene with a C-terminal His-tag. Following inoculation at 37 °C, the cells were incubated in self-inducible media (38) for 18 hours (shaking at 220 RPM, 16 °C). Centrifugation (3000 RPM, 30 min, 4 °C) was used to harvest the cells. Subsequently the cells were resuspended in TBS buffer (25 mM Tris + 2.7 mM KCl + 137 mM NaCl, pH 7.4; Alfa Aesar / Thermo Fisher Scientific, Cat # J60764) containing cOmplete protease inhibitor cocktail (Sigma Millipore) and lysed using sonication and pressurized cell disruption. Cell lysate was cleared by centrifugation at 12000 RPM for 1 hour at 4 °C. cOmplete His-Tag Purification Resin (Sigma Millipore) was applied for affinity purification. An additional wash step with 100 ml of detergent-containing buffer (25 mM Tris + 500 mM NaCl + 0.5 % N-Dodecyl-β-D-maltoside, pH 7.4) was introduced to remove endotoxin from the samples used for immunogen preparation. Samples were eluted using high imidazole buffer (25 mM Tris + 500 mM NaCl + 500 mM imidazole, pH 7.4). Proteins were then concentrated and buffer-exchanged to SEC buffer (25 mM Tris + 500 mM NaCl + 250 mM l-arginine + 5 % glycerol, pH 7.4) using Amicon ultrafiltration units, 10 kDa cutoff (Millipore Sigma). HiLoad 16/600 Superdex S200 pg column was used for final size exclusion purification step.

### Nanoparticle assembly studies

Three assembly reactions containing 5 µg of appropriate antigen-bearing component and equimolar amounts of corresponding assembly component were incubated at different temperatures (4, 25 and 37 °C) for 24 hours. Samples were run on NativePAGE 3-12% BisTris Protein Gels using the dark blue cathode protocol for the NativePAGE™ Novex® Bis-Tris Gel system (Thermo Fisher Scientific). Gels were fixed, de-stained as recommended by the protocol and imaged.

### Differential Scanning Fluorimetry

Measurements were performed on a Prometheus NT.48 NanoDSF instrument (NanoTemper Technologies) as described previously (72). Protein and nanoparticle samples were diluted to 0.5 mg/ml in the SEC buffer and loaded into NanoDSF capillaries (in triplicates). Tm measurement range was 20 – 95 °C at a rate of 1 °C/min. The first derivative curve was calculated from the raw data using the instrument software and the location of the maximum recorded as the Tm value.

### Negative stain electron microscopy

Negative stain electron microscopy experiments were performed as described previously (72, 73). Free nanoparticle components and purified nanoparticles were diluted to 20-50 µg/ml and loaded onto the carbon-coated 400-mesh Cu grid (glow-discharged at 15 mA for 25 s) for 10 s. After the sample was blotted off the grids were negatively stained with 2 % (w/v) uranyl-formate for 60 s. Data was collected on a Tecnai Spirit electron microscope, operating at 120 keV. Nominal magnification was set to 52,000 X with a pixel size of 2.05 Å at the specimen plane. Electron dose was adjusted to 25 e-/Å^2^ and the defocus was set at -1.50 µm. All micrographs were recorded on a Tietz 4k x 4k TemCam-F416 CMOS camera using Leginon automated imaging interface. Initial data processing was performed using the Appion data processing suite. For nanoparticle samples, approximately ∼1,000 particles were manually picked from the micrographs and 2D-classified using the Iterative MSA/MRA algorithm. For antigen-bearing components components and other trimer samples, 20,000 – 40,000 particles were auto-picked and 2D-classified using the Iterative MSA/MRA algorithm. For 3D classification and refinement, processing was continued in Relion/3.0 (74). Maps were segmented and color-coded in UCSF Chimera 1.13 (75).

### Cryo-EM grid preparation

Grids were prepared on a Vitrobot mark IV (Thermo Fisher Scientific). The setting were as follows: Temperature set to 10 °C, Humidity at 100 %, Blotting time varied in the 3-7 s range, Blotting force set to 0, Wait time of 10s. BG505-SOSIP-T33_dn10 nanoparticle sample was concentrated to 1.6 mg/ml and BG505-SOSIP-I53_dn5 nanoparticle was concentrated to 1.7 mg/ml. Lauryl maltose neopentyl glycol (LMNG) at a final concentration of 0.005 mM was used for sample freezing. Quantifoil R 2/1 holey carbon copper grid (Cu 400 mesh) were treated with Ar/O2 plasma (Solarus plasma cleaner, Gatan) for 10s before sample loading. Sample was mixed with the appropriate volume of LMNG solution and 3 µl immediately loaded onto the grid. Following the blot step the grids were plunge-frozen into nitrogen-cooled liquid ethane.

### Cryo-EM data collection

Cryo-grids were loaded into a Talos Arctica TEM (Thermo Fisher Scientific) operating at 200 kV, equipped with the K2 direct electron detector camera (Gatan) and sample autoloader. Total exposure was split into 250 ms frames with a total cumulative dose of ∼50 e^-^/Å^2^. Exposure magnification of 36,000 was set with the resulting pixel size of 1.15 Å at the specimen plane. For BG505-SOSIP-T33_dn10 nanoparticle imaging the nominal defocus range was -0.6 to -2.0 µm. The range was -0.8 to -2.0 for BG505-SOSIP-I53_dn5. Automated data collection was performed using Leginon software (76). The data collection details for the acquired datasets are presented in Supplement Table II.

### Cryo-EM image processing

MotionCor2 (77) was used to align and dose-weight the movie micrographs and the aligned micrographs were uploaded to cryoSPARC 2.9.0 (78). GCTF was then applied to estimate the CTF parameters. Particles were picked using template picker, extracted and 2D classified. Selected subsets of particles were then transferred to Relion/3.0 (74) for further processing. A reference model was generated using Ab-Initio Reconstruction in cryoSPARC 2.9.0. Multiple rounds of 3D classification and refinement were used to sort out a subpopulation of particles that went into the final 3D reconstructions. Tetrahedral and Icosahedral symmetry restraints were applied for all 3D refinement / classification steps during the processing of BG505-SOSIP-T33_dn10 and BG505-SOSIP-I53_dn5 datasets, respectively. A soft solvent mask around the nanoparticle core was introduced during the final 3D classification, refinement and post-processing steps in order to eliminate the signal originating from flexibly-linked BG505-SOSIP.v5.2(7S) trimers. Localized reconstruction v1.2.0 (64) was applied to obtain higher resolution information on the presented antigens. Vectors used for subparticle extraction were defined using sets of Chimera marker coordinates for each geometry (tetrahedral and icosahedral). Part of the signal corresponding to the nanoparticle core was subtracted from aligned particles and the trimer subparticles are extracted. Trimer subparticles are then subjected to 2D and 3D classification using a combination of Relion 3.0 and cryoSPARC 2.9.0 packages. Final subset of clean trimer subparticles was 3D-refined with C3 symmetry. A soft solvent mask around the reconstructed trimer was applied for refinement and post-processing steps. A graphical summary of the data processing approach and relevant statistics are displayed in Supplementary Figures S5 and S6. Final post-processed maps, half-maps and masks used for refinement and postprocessing were submitted to The Electron Microscopy Data Bank (EMDB). EMDB IDs: 21183 (I53_dn5 nanoparticle core), 21184 (BG505-SOSIP reconstructed from BG505-SOSIP_I53_dn5), 21185 (T33_dn10 nanoparticle core), 21186 (BG505-SOSIP reconstructed from BG505-SOSIP_T33_dn10).

### Model building and refinement

B-factor-sharpened maps corresponding to the T33_dn10 nanoparticle core and fused BG505-SOSIP.v5.2(7S) antigen generated in the previous step were used for model building and refinement. T33_dn10 model from Rosetta design was used for NP core refinement (with tetrahedral symmetry). BG505-SOSIP structure from PDB entry 5CEZ (65) was used as a starting model for trimer refinement (with C3 symmetry imposed). Iterative rounds of Rosetta relaxed refinement (62) and manual refinement in Coot (63) were performed to generate the final structures. EMRinger (79) and MolProbity (80) analysis was applied to evaluate the refined models. The refined models of T33_dn10 nanoparticle core and BG505-SOSIP (reconstructed from BG505-SOSIP-T33_dn10 nanoparticle) were submitted to the Protein Data Bank (PDB). PDB IDs: 6VFK (T33_dn10 nanoparticle core); 6VFL (BG505-SOSIP reconstructed from BG505-SOSIP_T33_dn10).

### Biolayer interferometry

Antibodies (IgG) were diluted in kinetics buffer (DPBS + 0.1 % BSA + 0.02 % Tween-20) to 5 µg/ml. Trimer and nanoparticle concentrations were normalized based on the molar concentration of the antigen (BG505-SOSIP.v5.2(7S) trimer) in each sample. Final BG505-SOSIP.v5.2(7S) concentration in the test samples was 500 nM. Free BG505-SOSIP.v5.2(7S) trimer was used as a positive control and a reference. Data was acquired on an Octet Red96 instrument (ForteBio). Antibodies were loaded onto anti-human IgG Fc capture (AHC) biosensors (ForteBio) and moved into the sample solutions at appropriate concentrations.

Association and dissociation steps were monitored for 180 s and 300 s, respectively. All data was analyzed using the ForteBio data processing package. Background was corrected by subtracting kinetics buffer dataset (negative control). The resulting binding curves for each antibody were corrected by aligning y-axes to the baseline step immediately preceding the association step and subsequently applying interstep correction between the association and dissociation. Baseline-corrected, aligned binding data was exported to Excel and plotted.

### Surface Plasmon Resonance

Antigenicity of ConM-SOSIP-T33_dn2 nanoparticles and free ConM-SOSIP.v7 envelope trimer was investigated by using surface plasmon resonance (SPR). All experiments were conducted at 25 °C on BIAcore 3000 instrument. HBS-EP (GE healthcare Life sciences) was used as running buffer throughout the analysis. To analyze binding of ConM-SOSIP-T33_dn2 nanoparticles and ConM-SOSIP.v7 trimers in solution, monoclonal antibodies (mAbs) were immobilized on CM3 sensor surface via covalently linked anti-Fc fragment. Affinity-purified goat anti-human IgG Fc (Bethyl Laboratories, Inc.) and goat anti-Rabbit IgG Fc (Abcam, USA) was amine coupled to CM3 surface, to capture human/Macaques and Rabbit mAbs, respectively. Both IgG Fc fragments were captured to a density of 5000 RU, as described elsewhere (37). ConM-SOSIP-T33_dn2 (5nM), and ConM-SOSIP.v7 trimer (20 nM) were used to analyze their binding to selected mAbs. In each experiment, 3 flow cells were used to capture IgG of mAbs on anti-Fc surface at an average density of 319 RU, with a standard deviation of ± 16 RU (SEM=1.5 RU), while one flow cell was used as reference. In each cycle, analyte (NP or trimer) was allowed to associate for 300 s, followed by dissociation for 600 s. At the end of each cycle, surface was regenerated with a single pulse of 75 µl of 10 mM Glycine (pH 2.0) at flow rate of 75 µl/min. MW of the peptide portion of ConM-SOSIP.v7 is 325 kDa with glycans and of ConM-SOSIP-T33_dn2 is 1.658 MDa with glycans. Data analysis and interpretation in experiments with nanoparticles and soluble antigens is described elsewhere (37).

### Nanoparticle stability tests

For pH sensitivity assessment, 3 aliquots of 5 µg of each nanoparticle sample (stored in TBS) were diluted using a set of concentrated Tris/Acetate buffers of different pH (100 mM Tris-Base + 150mM NaCl, pH adjusted with glacial acetic acid to pH 5, 7 and 9) and incubated at room temperature for 1 hour. Salt sensitivity assays were performed by diluting the same amount of each nanoparticle sample into buffers of different NaCl concentration (25 mM Tris + 25 / 150 / 1000 mM NaCl, pH 7.4), and subsequent incubation for 1 hour at room temperature. Temperature sensitivity was probed by incubating nanoparticles in TBS at a range of temperatures (25, 37, 50, 65 °C) for 1 hour. Sensitivity to freeze-thaw was probed by 1 or 2 rounds of flash-freezing in liquid nitrogen for 5 min followed by a gentle thaw at 4 °C for 1 hour. Sample homogeneity was assessed using Native PAGE (NativePAGE 3-12 % BisTris Protein Gels), dark blue cathode protocol for NativePAGE™ Novex® Bis-Tris Gel system (Thermo Fisher Scientific). Gels were fixed and de-stained using the recommended protocol and imaged.

### N-glycan analysis using hydrophilic interaction chromatography-ultra-high-performance liquid chromatography (HILIC-UPLC)

N-glycan profiling using HILIC-UPLC has been described in detail elsewhere (71, 81). In short, N-linked glycans were released from gp140 in-gel using PNGase F (New England Biolabs). The released glycans were subsequently fluorescently labelled with procainamide and excess label and PNGase F was removed using Spe-ed Amide-2 cartridges (Applied Separations). Glycans were analyzed on a Waters Acquity H-Class UPLC instrument with a Glycan BEH Amide column (2.1 mm x 100 mm, 1.7 μM, Waters). Fluorescence was measured, and data were processed using Empower 3 software (Waters, Manchester, UK). The relative abundance of oligomannose glycans was measured by digestion with Endoglycosidase H (Endo H; New England Biolabs). Digested glycans were cleaned using a PVDF protein-binding membrane (Millipore) and analyzed as described above.

### Site-specific glycan analysis using mass spectrometry

Env proteins were denatured for 1h in 50 mM Tris/HCl, pH 8.0 containing 6 M of urea and 5 mM dithiothreitol (DTT). Next, the Env proteins were reduced and alkylated by adding 20 mM iodacetamide (IAA) and incubated for 1h in the dark, followed by a 1h incubation with 20 mM DTT to eliminate residual IAA. The alkylated Env proteins were buffer-exchanged into 50 mM Tris/HCl, pH 8.0 using Vivaspin columns (3 kDa) and digested separately O/N using trypsin or chymotrypsin (Mass Spectrometry Grade, Promega) at a ratio of 1:30 (w/w). The next day, the peptides were dried and extracted using C18 Zip-tip (MerckMilipore). The peptides were dried again, re-suspended in 0.1 % formic acid and analyzed by nanoLC-ESI MS with an Easy-nLC 1200 (Thermo Fisher Scientific) system coupled to a Fusion mass spectrometer (Thermo Fisher Scientific) using higher energy collision-induced dissociation (HCD) fragmentation. Peptides were separated using an EasySpray PepMap RSLC C18 column (75 µm x 75 cm). The LC conditions were as follows: 275-minute linear gradient consisting of 0-32 % acetonitrile in 0.1 % formic acid over 240 minutes followed by 35 minutes of 80 % acetonitrile in 0.1 % formic acid. The flow rate was set to 200 nL/min. The spray voltage was set to 2.7 kV and the temperature of the heated capillary was set to 40 °C. The ion transfer tube temperature was set to 275 °C. The scan range was 400-1600 m/z. The HCD collision energy was set to 50 %, appropriate for fragmentation of glycopeptide ions. Precursor and fragment detection were performed using an Orbitrap at a resolution MS1= 100,000. MS2= 30,000. The AGC target for MS1=4e^5^ and MS2=5e^4^ and injection time: MS1 = 50 ms MS2 = 54 ms.

Glycopeptide fragmentation data were extracted from the raw file using Byonic^TM^ (Version 3.5) and Byologic^TM^ software (Version 3.5; Protein Metrics Inc.). The glycopeptide fragmentation data were evaluated manually for each glycopeptide; the peptide was scored as true-positive when the correct b and y fragment ions were observed along with oxonium ions corresponding to the glycan identified. The MS data was searched using a standard library for HEK293F expressed BG505 SOSIP.664. The relative amounts of each glycan at each site as well as the unoccupied proportion were determined by comparing the extracted chromatographic areas for different glycotypes with an identical peptide sequence. The precursor mass tolerance was set at 4 ppm and 10 ppm for fragments. A 1 % false discovery rate (FDR) was applied. The relative amounts of each glycan at each site as well as the unoccupied proportion were determined by comparing the extracted ion chromatographic areas for different glycopeptides with an identical peptide sequence.

### Site-specific analysis of low abundance N-glycan sites using mass spectrometry

To obtain data for sites that frequently present low intensity glycopeptide the glycans present on the glycopeptides were homogenized to boost the intensity of these peptides. A separate tryptic digest was used for this workflow. This analysis loses fine processing information but enables the ratio of oligomannose: complex: unoccupied to be determined. The peptides were first digested with Endo H (New England Biolabs) to deplete oligomannose- and hybrid-type glycans and leave a single GlcNAc residue at the corresponding site. The reaction mixture was then dried completely and resuspended in a mixture containing 50 mM ammonium bicarbonate and PNGase F (New England Biolabs) using only H2^18^O (Sigma-Aldrich) throughout. This second reaction cleaves the remaining complex-type glycans but leaves the GlcNAc residues remaining after Endo H cleavage intact. The use of H2^18^O in this reaction enables complex glycan sites to be differentiated from unoccupied glycan sites as the hydrolysis of the glycosidic bond by PNGaseF leaves a heavy oxygen isotope on the resulting aspartic acid residue. The resultant peptides were purified as outlined above and subjected to reverse-phase (RP) nanoLC-MS. Instead of the extensive N-glycan library used above, two modifications were searched for: +203 Da corresponding to a single GlcNAc, a remnant of an oligomannose/hybrid glycan, and +3 Da corresponding to the ^18^O deamidation product of a complex glycan. A lower HCD energy of 27 % was used as glycan fragmentation was not required. Data analysis was performed as above and the relative amounts of each glycoform determined, including unoccupied peptides.

### B cell activation assays

B cell activation experiments were performed as previously described (55). K46 cells expressing doxycycline-inducible VRC01, PGT145 and PGT121 receptors (in a form of IgM) were grown in advanced Dulbecco’s modified Eagle’s medium (DMEM) (Gibco), supplemented with 10 % fetal calf serum, penicillin/streptomycin antibiotics, and puromycin (2 μg/ml; Gibco). 1 µg/ml Doxycycline was added overnight to induce BCR expression. Cells were spun down (500 g, 3 min, room temperature) and resuspended in RPMI 1640 media supplemented with GlutaMAX (1X), 10% FBS, penicillin/streptomycin antibiotics (1X), 2-mercaptoethanol, with 1.5 µM Indo-I fluorescent dye (Thermo Fisher Scientific). Cells were incubated for 1 hour with the dye at 37 °C, washed 3 times with cold PBS and transferred to fresh cold media that does not contain Indo-I (cell density: 1 * 10^6^ cells/ml). Calcium mobilization was recorder on an LSR II flow cytometer (BD Biosciences) by measuring the 405/485-nm emission ratio of Indo-1 fluorescence upon UV excitation at room temperature. Trimer and nanoparticle samples were normalized to contain an equimolar amount of BG505-SOSIP.v5.2(7S) antigen in each sample (7.5 nM final assay concentration). Anti-mouse IgM antibody (Jackson ImmunoResearch) at 10 µg/ml was used as a positive control. 1 ml Aliquots of Indo-I treated cells (cell density: 1 * 10^6^ cells/ml) were incubated for 60 s for the baseline signal to be recorded, after which they are stimulated with an antigen for 180 s. Immediately following this step, ionomycin (at 1 µg/ml) is added to the cells and the signal was recorded for another 60 s to verify Indo-I loading. Data analysis was performed using FlowJo (Tree Star, Ashland, OR).

### Immunization experiments

Immunizations and blood draws were performed by Covance Research Products Inc. under permits with approval number C0171-017 (Denver, PA, USA). 2 groups of 5 female New Zealand White Rabbits were immunized at weeks 0, 4 and 20 with ConM-SOSIP.v7 (Group 1, 30 µg/dose) and ConM-SOSIP-T33_dn2 nanoparticle (Group 2, 43 µg/dose) Total dose was normalized to achieve an equivalent molar concentration of ConM-SOSIP.v7 antigen across all animals. Peptidic molecular weight of the protein was used for dose calculations (disregarding glycan). Immunogens were formulated with GLA-LSQ adjuvant (25 µg GLA and 10 µg QS21 per dose; The Infectious Disease Research Institute (IDRI), Seattle, WA). Rabbits were immunized with antigen-adjuvant formulation via intramuscular route. The dose was split in half and injected into both quadriceps. Blood draws were performed at weeks 0, 2, 4, 6, 8, 12, 16, 20 and 22. These experiments were performed in parallel with ConM-SOSIP.v7 immunization experiments reported elsewhere (37) and the data acquired for ConM-SOSIP.v7 control group (Group 1) is shared between two studies.

### ELISA binding assays

Experiments were performed as described previously (37). ConM-SOSIP.v7 carrying a C-terminal His-tag (diluted to 6.5nM in TBS) was immobilized onto 96-well Ni-NTA ELISA plates (Qiagen) by incubation for 2 hours at room temperature after which the plates were washed 3 times with TBS. Plates were blocked with TBS + 2 % skimmed milk and washed 3 times with TBS. For experiments with T33_dn2 nanoparticle core, 6nM final concentration of the antigen was used for plate preparation. Serial three-fold sera dilutions were prepared in the binding buffer (TBS + 2 % skimmed milk + 20 % sheep serum) starting at a minimum of 1:200. For experiments with T33_dn2 nanoparticle core the starting dilution was 1:1000. Diluted samples were incubated on the plates for 2 hours at room temperature. Following 3 washes with TBS, HRP-conjugated goat anti-rabbit IgG (Jackson Immunoresearch) in TBS + 2% skimmed milk was added for 1 hour at room temperature. Detection antibody was diluted 1:3000. Plates were washed 5 times with TBS + 0.05% Tween-20, followed by the addition of developing solution (1% 3,3’,5,5’-tetranethylbenzidine (Sigma-Aldrich) + 0.01% hydrogen peroxide, 100 mM sodium acetate and 100 mM citric acid) was added. Colorimetric endpoint development was allowed to proceed for 3 min before termination by 0.8 M H2SO4. Endpoint titers were determined using Graphpad Prism software.

### Virus neutralization assays

Pseudovirus neutralization assays were performed at Amsterdam University Medical Center (AUMC) as described previously (82). Serial 3-fold dilutions of rabbit sera samples (starting at 1:20 or 1:100 dilution) were prepared and tested against ConM-pseudotyped virus in TZM-bl cells in a 96-well format. Midpoint titers (IC50) were determined using Graphpad Prism software.

### EM-based polyclonal epitope mapping – Preparation of Fab and complex samples

Experiments were performed as described previously (69). Briefly, serum samples (weeks 4 and 22) from 2 immunized animals with highest neutralization titers in each group were selected for polyclonal epitope mapping. Animal ID of selected animals: r2381 (Group 1), r2382 (Group 1), r2383 (Group 2) and r2385 (Group 2). IgGs were purified from ∼1-3 ml of serum using equal volume of settled Protein A Sepharose resin (GE Healthcare). Samples were eluted off the resin with 0.1 M glycine pH 2.5 and immediately neutralized with 1 M Tris-HCl pH 8. Amicon ultrafiltration units, 10 kDa cutoff (Millipore Sigma) were used to concentrate the purified IgG and buffer exchange to the digestion buffer (PBS + 10 mM EDTA + 20 mM cysteine, pH 7.4). IgG samples were digested for 5 hours at 37 °C using 50 µl of settled papain-agarose resin (Thermo Fisher Scientific). Fc and non-digested IgG were removed by 1-hour incubation with Protein A Sepharose resin, using 0.2ml packed resin per 1 mg of starting IgG amount (room temperature). Fab samples were concentrated to ∼2-3 mg/ml using Amicon ultrafiltration units, 10 kDa cutoff (Millipore Sigma), and in the process buffer was exchanged to TBS. Final Fab yields were ∼300µg. Initial assembly trials were performed with 250 µg of purified Fab samples and 15 µg of ConM-SOSIP.v7 for ∼18 hours at room temperature, but we observed that Fab binding caused >70% of these trimers to dissociate into gp120-gp41 monomers. Accordingly, it was not possible to use the EMPEM data to reliably assign epitopes (Supplementary Figure S9.a). Instead, we complexed the purified Fabs with ConM-SOSIP.v9 trimers, which contain additional stabilizing mutations that do not affect antigenicity (see Methods, and Supplementary Table I, Supplementary Figure S9. a and b). When the more stabilized trimer was used for EMPEM, only <5% of the trimers dissociated into gp120-gp41 monomers after Fab binding, which allowed us to assign the recognized epitopes more reliably (Supplementary Figure S9.a). Hence, the complexing was performed with 250 µg of polyclonal Fab and 15 µg of ConM-SOSIP.v9. Complexes were purified using SEC (Superose 6 Increase column) with TBS as a running buffer, concentrated with Amicon ultrafiltration units (10 kDa cutoff) and immediately loaded onto the carbon-coated 400-mesh Cu grid (glow-discharged at 15 mA for 25 s). Samples were diluted in TBS to 50 µg /ml prior to loading. Grids stained with 2 % (w/v) uranyl-formate for 60 s.

### EM-based polyclonal epitope mapping – Sample imaging and data processing

Imaging was performed as described above in the negative stain electron microscopy method section. All initial processing was performed using the Appion data processing package (83). Approximately, 120,000 – 150,000 particles are picked and extracted. Particles were then 2D-classified in Relion 3.0 (74) into 250 classes (50 iterations), and particles with complex-like features (∼70-90% of the starting number) were selected for 3D sorting in Relion 3.0. A low-resolution model of non-liganded HIV Env ectodomain was used as a reference for all 3D steps. Initial 3D classification was performed with 40 classes. Particles from similar looking classes were then pooled and reclassified. A subset of 3D classes with unique structural features (in terms of Fab specificities) was subjected to 3D auto-refinement in Relion 3.0. Maps from 3D refinement were loaded into UCSF Chimera 1.13 (75) for visualization, segmentation and figure preparation. 3D refinement was performed on a subset of 2D-cleaned particles (following the initial 2D classification step and before any 3D classification) and the refined model is submitted to EMDB. The list of EMDB IDs: 21175 (ConM-SOSIP.v9 + Wk4-r2381 polyclonal Fab); 21176 (ConM-SOSIP.v9 + Wk4-r2382 polyclonal Fab); 21177 (ConM-SOSIP.v9 + Wk4-r2383 polyclonal Fab); 21178 (ConM-SOSIP.v9 + Wk4-r2385 polyclonal Fab); 21179 (ConM-SOSIP.v9 + Wk22-r2381 polyclonal Fab); 21180 (ConM-SOSIP.v9 + Wk22-r2382 polyclonal Fab); 21181 (ConM-SOSIP.v9 + Wk22-r2383 polyclonal Fab), 21182 (ConM-SOSIP.v9 + Wk22-r2385 polyclonal Fab). Full particle stacks and 3D models used for Fab segmentation and generation of composite figures are available upon request.

## Acknowledgements

The authors thank Bill Anderson, Hannah L Turner and Charles A Bowman for their help with electron microscopy, data acquisition and processing. The authors also thank Kimmo Rantalainen for providing a small-scale screening protocol for expression and evaluation of fusion constructs. The authors thank Lauren Holden for her help on the preparation of this manuscript. This work was supported by grants from the National Institute of Allergy and Infectious Diseases Center for HIV/AIDS Vaccine Immunology and Immunogen Discovery UM1AI100663 (M.C., D.N. A.B.W.), Center for HIV/AIDS Vaccine Development UM1AI144462 (M.C., D.N. A.B.W.), P01 AI110657 (J.P.M., R.W.S., and A.B.W.), R01 AI36082 (J.P.M.), R01AI073148 (D.N.) and R01AI128836 (D.N.); and by the Bill and Melinda Gates Foundation and the Collaboration for AIDS Vaccine Discovery (CAVD) OPP1156262 (N.P.K., D.B.), OPP1120319 (N.P.K., D.B.), OPP1115782 (A.B.W.), OPP1111923 (D.B., N.P.K., J.P.M., and R.W.S.), OPP1132237 (D.B., N.P.K., J.P.M., and R.W.S.) and OPP1196345 (D.N. and A.B.W.); and by the National Science Foundation grant DMREF 1629214 (N.P.K., D.B.); and by the Howard Hughes Medical Institute (D.B.). This work was also supported by the European Union’s Horizon 2020 research and innovation program under grant agreement No. 681137 (M.C., R.W.S.). C.A.C. is supported by a NIH F31 Ruth L. Kirschstein Predoctoral Award Al131873 and by the Achievement Rewards for College Scientists Foundation. This work was partially funded by IAVI with the generous support of USAID, Ministry of Foreign Affairs of the Netherlands, and the Bill & Melinda Gates Foundation; a full list of IAVI donors is available at www.iavi.org. The contents of this manuscript are the responsibility of the authors and do not necessarily reflect the views of USAID or the US Government.

## Supporting Information

**Supplementary Table I.** Protein sequences of constructs used in this study

**Supplementary Table II.** Cryo EM data collection statistics

**Supplementary Table III.** Model building and refinement statistics for T33_dn10 nanoparticle core and presented BG505-SOSIP.

**Supplementary Table IV.** Anti-trimer binding antibody titers against ConM-SOSIP.v7

**Supplementary Table V.** Autologous neutralization titers against ConM-based pseudovirus

**Supplementary Figure S1. Nanoparticle library evaluated in this study.** (a) Structural models of nanoparticle candidates derived from Rosetta_design. For clarity, trimeric antigen-bearing component is shown in orange and assembly component in blue. (b) Geometric properties of different nanoparticle candidates.

**Supplementary Figure S2. Purification and characterization of different antigen-presenting components and assembled nanoparticles.** (a) SEC curves of BG505-SOSIP.v5.2(7S) and BG505-SOSIP-fused nanoparticle components. (b) SDS PAGE analysis of the purified assembly component for T33_dn2, T33_dn10 and I53_dn5 nanoparticle systems. (c) SEC purification of different nanoparticle candidates after assembly. (d) SDS PAGE gel of the purified nanoparticles confirming the presence of both, antigen-bearing and assembly components.

**Supplementary Figure S3. Site specific glycan analysis of BG505-SOSIP-bearing components and free BG505-SOSIPv5.2(7S).** The table shows the glycoforms found at each potential N-linked glycosylation site (PNGS), compositions corresponding to oligomannose/hybrid-type glycans are colored green and fully processed complex type glycans are colored magenta. PNGS with no attached glycan are colored grey. Oligomannose-type glycans are categorized according to the number of mannose residues present, hybrids are categorized according to the presence/absence of fucose and complex-type glycans are categorized according to the number of processed antenna and the presence/absence of fucose. Sites that could only be obtained from low intensity peptides cannot be distinguished into the categories in the table and so are merged to cover all oligomannose/hybrid compositions or complex-type glycans

**Supplementary Figure S4. Nanoparticle stability studies.** Native PAGE assays were used for evaluation of nanoparticle integrity following the incubation under the specified conditions.

**Supplementary Figure S5. Cryo-EM analysis of BG505-SOSIP-T33_dn10 nanoparticle.** Schematic representation of the data processing workflow with relevant statistics.

**Supplementary Figure S6. Cryo-EM analysis of BG505-SOSIP-I53_dn5 nanoparticle.** Schematic representation of the data processing workflow with relevant statistics.

**Supplementary Figure S7. ConM-SOSIP-T33_dn2 nanoparticle purification and characterization.** (a) SEC purification of ConM-SOSIP-T33_dn2A and 2D class-averages from negative-stain-EM analysis. (b) SEC purification of assembled ConM-SOSIP-T33_dn2 and NS-EM analysis of the purified nanoparticles (raw micrograph and 2D class averages). (c) SPR-based characterization of the antigenicity of purified nanoparticles. ConM-SOSIP.v7 trimer was used as a reference. In addition to affinity, SPR signal is also a function of antigen size (molecular weight). MW of the ConM-SOSIP-T33_dn2 nanoparticle is ∼5.1 times higher than that of soluble ConM-SOSIP.v7 trimer. See methods section for data analysis information.

**Supplementary Figure S8. Extended immunization data.** (a) Anti-nanoparticle core response (ELISA binding titers) in individual Group 2 animals, with the mean value indicated by the solid line. The dashed line represents the assay detection limit. (b) Ratios of NAb titers and anti-trimer binding antibody titers in sera from individual animals in Group 1 (open circles) and Group 2 (black squares) were calculated at weeks 4, 8 and 22. Scatter plots are shown with mean values indicated. Red asterisks indicate sera samples where neutralization titers were below the level of detection (1:20 titer). An AUC statistical analysis of the titer ratio values for Group 1 versus Group 2 as a function of time results in p = 0.016.

**Supplementary Figure S9. Comparison of ConM-SOSIP.v7 and ConM-SOSIP.v9.** (a) EMPEM data derived using ConM-SOSIP.v7 (top) and ConM-SOSIP.v9 (bottom), complexed with polyclonal Fab sample isolated from the serum of rabbit r2381 (Grp1) at week 22 time point. Representative raw micrographs (left), 2D classes (middle) and 3D classes (right) are shown. Red circles mark the 2D classes of ConM-SOSIP.v7 gp120-gp41 monomers bound to one or more Fabs. Reconstructed monomer-like 3D classes are shown. Fabs are easily discernable in most 3D classes but epitope assignment is very challenging due to high degree of heterogeneity. Complexing with ConM-SOSIP.v9 results in significantly lower percentage of disassembled trimers (i.e. monomers), which can be observed in 2D and 3D classes. (b) Comparison of stabilizing mutations in different ConM-SOSIP constructs.

**Supplementary Figure S10. Extended data for EMPEM analysis of antibody responses in immunized rabbits.** Sample micrograph, 2D class averages and sample 3D reconstructions obtained for antibodies isolated from the specified rabbits at (a) week 4 and (b) week 22.

